# Clust: automatic extraction of optimal co-expressed gene clusters from gene expression data

**DOI:** 10.1101/221309

**Authors:** Basel Abu-Jamous, Steven Kelly

## Abstract

Identification of co-expressed gene clusters can provide evidence for genetic or physical interactions between genes. Thus, co-expression clustering is a routine step in large-scale analyses of gene expression data. We show that commonly used clustering methods produce results that substantially disagree with each other, and do not match the biological expectations of co-expressed gene clusters. Furthermore, these clusters can contain up to 50% unreliably assigned genes. Consequently, downstream analyses of these clusters (e.g. functional term enrichment analysis) suffer from high error rates. We present *clust*, an automated method that solves these problems by extracting clusters that match the biological expectations of co-expressed genes. Using 100 datasets from five model organisms we demonstrate that clusters generated by *clust* are better than those produced by other methods, both numerically and for use in functional analysis. Finally, we show that *clust* can simultaneously cluster multiple datasets, enabling users to leverage the large quantity of public expression data for novel comparative analysis.

## Introduction

Gene transcription is dynamically and coordinately regulated in all living organisms. Such coordinate regulation is manifest as concordant changes in the transcript abundance of genes in time series and perturbation-response datasets. Gene transcription is regulated by the binding of transcription factors to DNA/chromatin elements located in promoter or enhancer regions of genes. Typically, transcription factors comprise ∼10% of the total number of genes in a genome, and complex spatio-temporal patterns of transcription are achieved through the combinatorial action of these genes in regulatory networks ^1^. The combinatorial nature of these networks means that their behaviour is inherently conditional. That is, genes that appear co-expressed under one condition are not necessarily co-expressed under all conditions. A corollary of this is that within any one experimental context (e.g. time series spanning some biological process or perturbation-response experiment) not all genes will be behaving coordinately. Instead, subsets of genes have the right combination of regulators to behave coordinately during the experimental context while others are following patterns of regulation that are independent of the experimental design. Thus, within a given observation window (i.e. experimental context) it is not expected that all genes can be assigned to a limited set of coordinate behaviours ^2,3^.

Given that only subsets of genes are likely to be co-expressed within a particular context, it follows that identification of these subsets is a data extraction problem and not a data partitioning problem. That is, the aim is to identify and extract the cohorts of genes that are behaving coordinately from the complete set of genes that are detected within a particular context, and is not to partition the complete set of genes into a set of gene clusters. In practice, clustering methods have been widely applied to gene expression data with the expectation that they will identify the complete set of discrete cohorts of genes that have co-ordinated behaviours (i.e. the clusters of co-expressed genes), and that all of genes that exhibit those behaviours will be assigned to the correct cluster. However, the vast majority of methods that aim to identify cohorts of co-expressed genes are based on data partitioning (e.g. Markov clustering ^4^, k-means ^5^, hierarchical clustering ^6^, and self-organising maps ^7^). These approaches attempt to assign all genes to a finite set of clusters, with the number of clusters determined by numerical optimisation of a data partitioning metric ^8^. Thus, genes that are not co-expressed in the context under investigation are also assigned to their “best-fitting” cluster such that the majority of clusters will contain both co-expressed and non-co-expressed genes. This result does not adhere to the expectation of the biological properties of a co-expressed gene cluster, i.e. that each cluster contains only those genes that exhibit co-ordinate behaviour in the experimental or biological context under question and that no two clusters should have an identical profile.

Here we show through analysis of 100 real biological datasets from five model organisms that application of data partitioning-based clustering methods to gene expression data generates clusters that include substantial numbers of unreliably assigned genes, i.e. genes that should have been excluded. Such unreliable content comprises up to about 50% of these clusters. To address this problem we provide a novel method called “*clust*” for cluster extraction from gene expression data. *Clust* is designed to extract co-expressed clusters of genes that satisfy the biological expectations of a co-expressed gene cluster. We show that *clust* satisfies these expectations by extracting co-expressed clusters with lower levels of dispersion than any data partitioning method. We also show that the clusters produced by *Clust* do not contain unreliably clustered genes typical of data partitioning methods. Furthermore, we show that the clusters extracted by *clust* are well enriched with functional terms, indicating their biological relevance. Finally, we demonstrate the ability of *clust* to extract clusters of consistently co-expressed genes in multiple datasets simultaneously, a feature that allows researchers with multiple datasets relating to the same biological question to analyse them collectively.

## Results

### Problem definition, aim and approach

Gene expression datasets (RNA-seq and microarray) contain quantitative estimates (observations) of mRNA abundance for a set of genes at multiple experimentally, spatially, or temporally discrete conditions. Across these conditions, it is expected that the mRNA abundance of transcriptionally co-regulated genes will exhibit coordinate behaviour. These co-regulated cohorts of genes include those that are inherent modules of the system being studied, as well as those that may be conditional on applied experimental perturbations. The observations also include transcript abundance estimates for genes that are behaving independently in the experimental series. Furthermore, for genes that are transcriptionally co-regulated, variance in RNA processing and mRNA half-life cause fluctuations in transcript abundance such that abundance estimates are inherently noisy. Thus, the goal of gene expression clustering is to identify and extract the discrete cohorts of genes whose transcripts are behaving coordinately (albeit with biological noise) across the observations under consideration.

Fig. 1 presents simulated gene expression data to illustrate the problem of extracting distinct cohorts of co-expressed genes. Each simulated dataset contains 500 genes, with 100 genes in each of three distinct clusters and 200 genes that do not belong to any cluster. Fig. 1a shows the same clusters simulated with increasing levels of biological noise (D1 to D4) and Fig. 1b shows the desired results. That is, to extract three distinct clusters of genes (C1 to C3) while discarding the genes that behave independently. In conflict with the desired goal, data partitioning methods require all genes to be included in one of the clusters. For example, application of *k*-means (the most commonly used method for analysing gene expression datasets) recovers the three simulated profiles. However, each cluster also contains a large cohort of genes that do not share the same expression profile. This inclusion results in clusters with high levels of dispersion and high levels of inter-cluster similarity, violating the expectations of co-expressed gene clusters, and producing clusters whose gene assignment is unreliable. *Clust* is designed to address this problem by extracting the largest and least dispersed set of clusters whose profiles are distinct and exclude those genes that do not belong to these clusters. That is, to identify and extract the complete set of genes that are exhibiting coordinate behaviour in the experimental series under consideration. The results of applying *clust* to these demonstrative datasets are included in Supplementary Figure S1.

**Figure 1.**
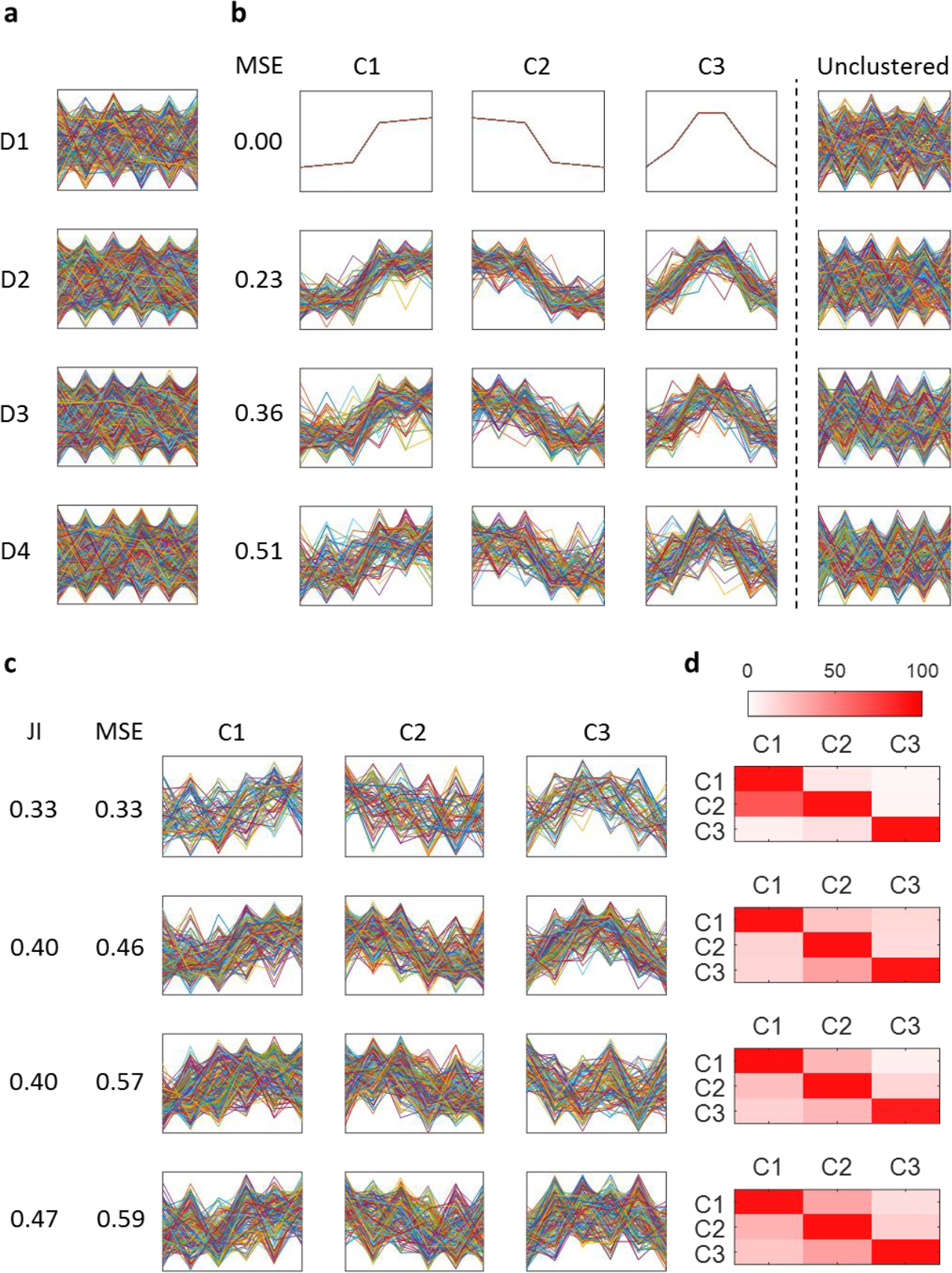
xpectations and outcomes for application of data-partitioning methods to coexpression clustering. (**a & b**) Simulated gene expression data for 500 genes with increasing noise (D1 – D4). (**a**) All genes. (**b**) profiles of the genes in each of the three simulated clusters as well as the extra unclustered genes at each one of the four levels of dispersion. The horizontal axis of each plot represents the six different conditions/samples, while the vertical axis represents gene expression values. (**c**) The results of applying a partitioning method (*k*-means in this case) to the same simulated datasets. (**d**) Heat-maps that show the percentage of genes in a cluster that also fit well within each one of the other clusters.

### Data sources and comparative methods

To demonstrate the performance characteristics of *clust* on real biological datasets, the method was applied to 100 different gene expression datasets (Supplementary Table S1). These datasets comprised ten microarray datasets and ten RNA-seq datasets from each of five different model organisms; *Homo sapiens, Mus musculus, Drosophila melanogaster, Arabidopsis thaliana*, and *Saccharomyces cerevisiae.* To put these performance characteristics in context, five of the most commonly used co-expression clustering methods (*k*-means, Markov clustering (MCL), hierarchical clustering (HC), WGCNA, and self-organising maps (SOMs)) were also applied to these datasets. For each of these additional methods, the best-practice operating procedures were followed as described in Methods.

### Clust robustly extracts tight and non-overlapping clusters

As *clust* is a cluster extraction method, it does not necessarily assign all genes to clusters. On average across the 100 test datasets *clust* assigned 50% of the input genes to clusters (Fig. 2a), and produced sets of clusters that have significantly lower dispersion than those produced by MCL (p-value 2.5×10^−31^), k-means (p-value 4.6×10^−7^), HC (p-value 2.4×10^−13^), WGCNA (p-value 7.7×10^−11^), or SOMs (p-value 4.8×10^−24^) (Fig. 2b). Clusters produced by *clust* are discrete, such that genes assigned to one cluster do not fit within the profile boundaries of any other cluster (JI = 0 for all clusters, Fig. 2c). This is not the case for data partitioning methods, where 10% to 50% of the genes that are included in a given cluster also fit within the boundaries of at least one other cluster (Fig. 2c). Thus, application of data partitioning methods to gene expression data produces clusters that are not discrete and contain between 10% and 50% unreliably assigned genes (Supplementary Table S2).

**Figure 2.**
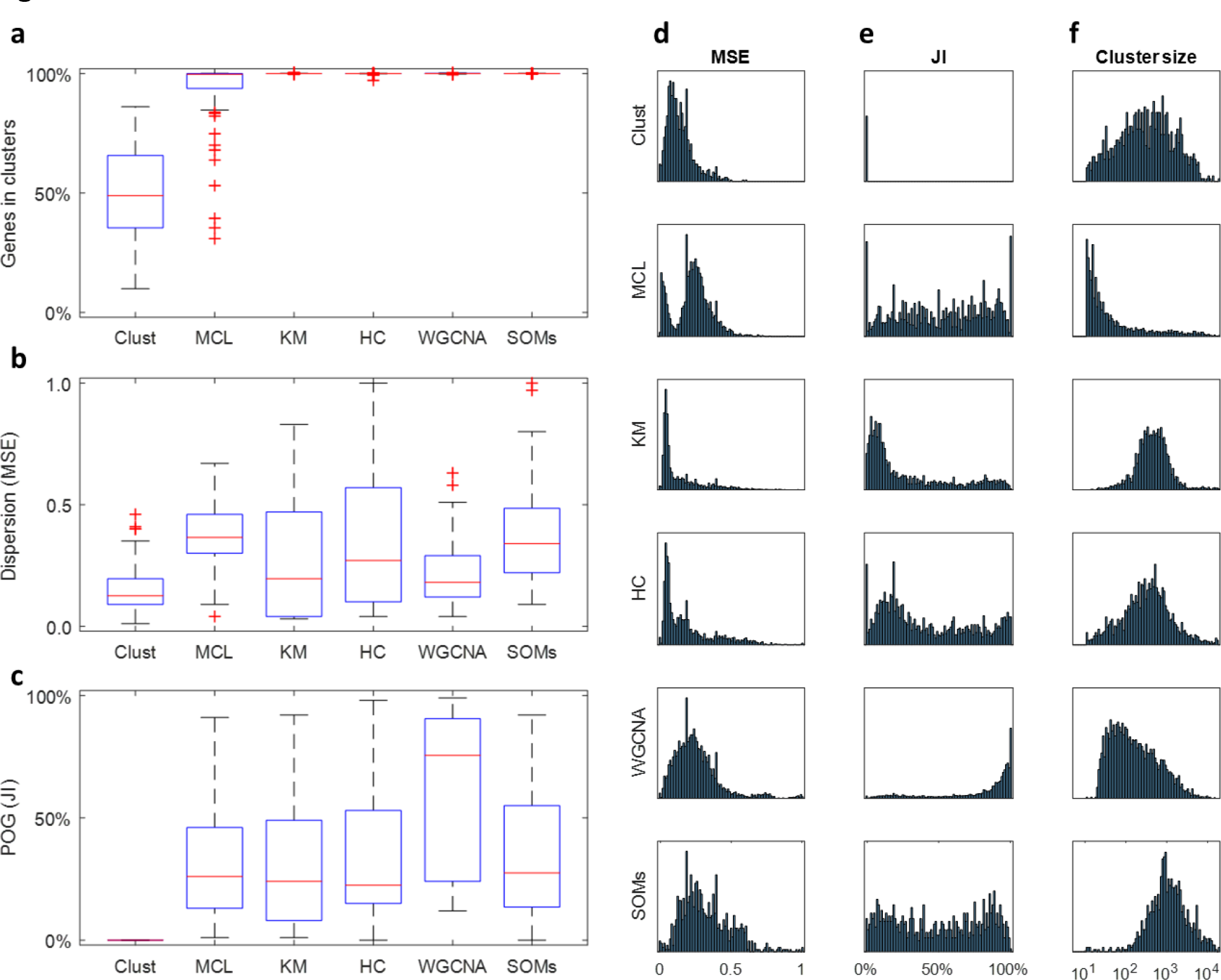
Evaluation of the performance of clustering methods. (**a-c**) Evaluation of clustering performance over all 100 datasets. (**a**) the percentage of input genes that were included in clusters; (**b**) the average dispersion of clusters measured by weighted-averaging of individual cluster MSE values; (**c**) percentage of the overlap amongst clusters, as measured by JI index. (**d-e**) Distributions of individual cluster properties for all 100 datasets. MSE values (**d**), JI values (**e**), and cluster sizes (**f**). (Supplemental Tables S2 and S3 and Supplemental Figures S3 to S10).

In addition to producing datasets that are both lower in dispersion and discrete, the distribution of the properties of *clust* clusters is also unimodal (Fig. 2d-f and Supplementary Table S3). In contrast, cluster dispersion is multimodal for MCL, *k*-means, HC, and SOMs (Fig. 2d); cluster overlap is multi-modal or uniformly distributed for MCL, KM, HC and SOMs (Fig. 2e); cluster size is biased towards small clusters for MCL and WCGNA (Fig. 2f). Thus, the individual clusters returned by data partitioning methods are inconsistent and vary considerably in their quality. In contrast, the quality of clusters produced by *clust* is unimodal such that all clusters can be considered to be drawn from a single population of clusters. This property of *clust* clusters is independent of the number of genes in a given cluster (Supplementary Figures S3 and S4). In contrast, the properties of clusters returned by data partitioning methods display a significant dependency on cluster size, such that larger clusters have both higher dispersion and a higher proportion of genes that fit within the boundaries of other clusters (Supplementary Figures S3 and S4).

Fig. 3 shows a comparative example of the clusters produced by each one of the six clustering methods when applied to one of the 100 datasets (D82, Supplementary Tables S1 and S2). This dataset was chosen as it is the one with the most similar results across all the tested methods and a similar figure showing the first up to 14 clusters produced by each method for all 100 datasets are provided for download from the Zenodo repository at 10.5281/zenodo.1169191. The reduced MSE and JI of *clust* in comparison to other methods is readily apparent from visual inspection of the gene expression profiles of genes assigned to each cluster in Fig. 3. Importantly, this is not at the expense of the clusters’ sizes. For instance, cluster C1 produced by *clust* is less dispersed and contains more genes than the most similar cluster generated by *k*-means, HC, or WGCNA (also labelled C1).

**Figure 3.**
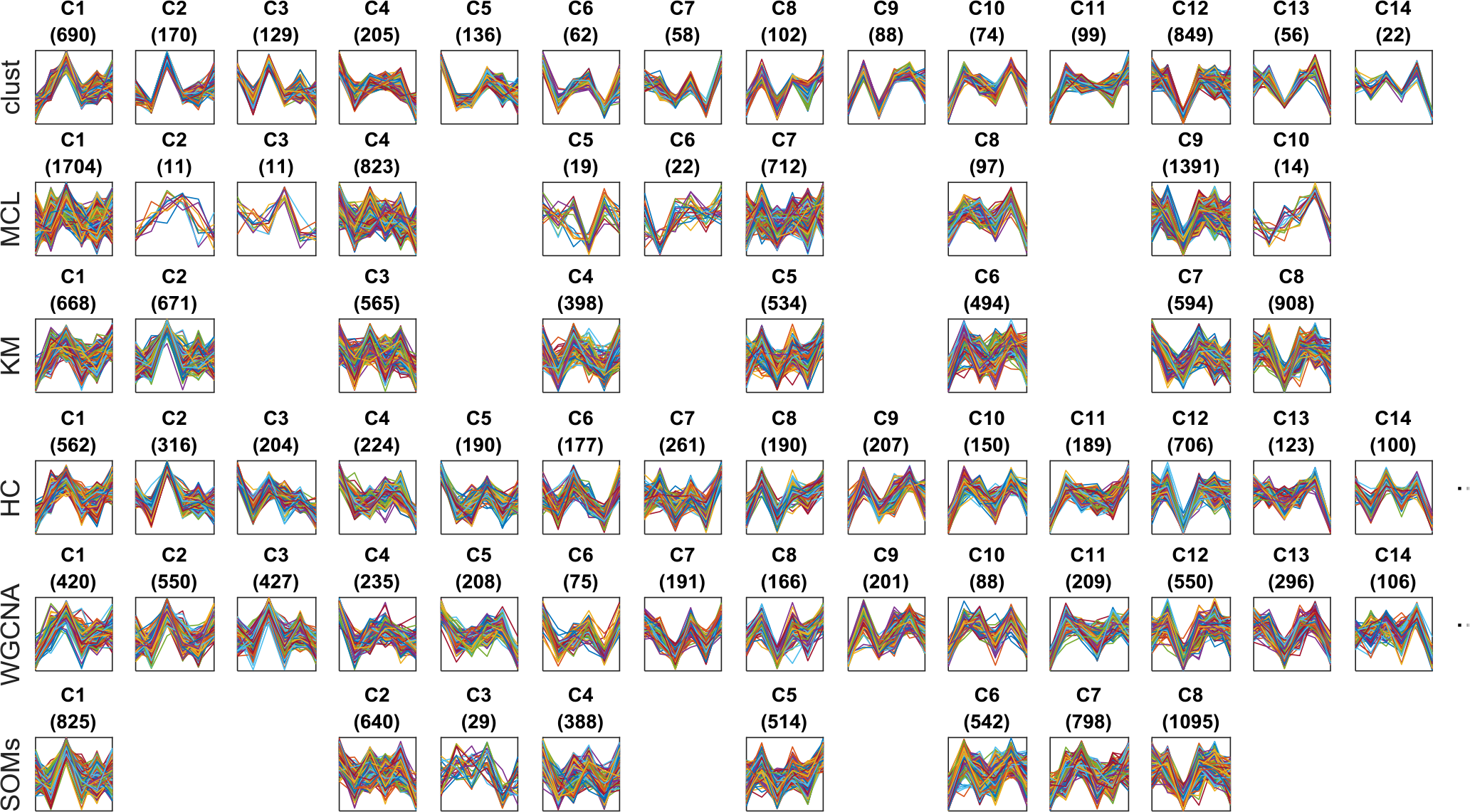
Profiles of the genes in the clusters generated by each method when applied to the dataset D82. This figure visually shows a sample of the results of each one of the methods when applied over the same dataset, which is the dataset D82 (Supplementary Tables S1 and S2). This dataset was chosen out of the 100 datasets because the numbers of clusters generated by the six methods are more similar to each other at this dataset than any other dataset (measured by the least squares metric). Datasets with less than five clusters generated by any method were avoided. The numbers of clusters generated for this dataset by clust, MCL, k-means, HC, WGCNA, and SOMs were 14, 10, 8, 23, 21, and 8, respectively. This figure shows all 14 clusters generated by clust in the first row. Then, the most similar clusters generated by the other methods to each one of the 14 clust’s clusters are aligned below them. When there are more than 14 clusters generated by the same method, only the 14 clusters which are most similar to the clust’s clusters are displayed here. The title of each sub-plot shows the name of the cluster and the number of genes in that cluster between parentheses. The horizontal axis of each subplot represents the six samples in the dataset D82 while the vertical axis represents the normalised gene expression value. The profiles of all individual genes in a cluster are drawn as lines on top of each other in its corresponding sub-plot.

None of the six methods, including *clust*, behaves differently on datasets from different species (Supplementary Figures S5 and S6). Therefore, the species from which the data was produced is not a factor that affects the performance of any of these clustering methods. However, the dispersion of clusters produced by *k*-means, HC, WGCNA, and SOMs, is dependent on the number conditions under consideration such that the more conditions being considered the worse the results of the clustering (Supplementary Figure S7). In contrast, the behaviour of *clust* is unaffected by the number of genes or the number of conditions that are under consideration (Supplementary Figures S7, S8, S9, and S10). Thus, unlike all of the other tested methods the behaviour of *clust* is consistent for different types or quantities or input data.

### Clusters extracted by clust have reliable enriched functional terms

One of the most commonly applied tests to co-expressed clusters of genes is functional term enrichment, as a cluster of co-expressed genes is expected to be enriched with genes that have related biological roles. As *clust* assigns on average 50% of genes to clusters, it was investigated how this reduction in gene number affects the detection of functional term enrichment. To do this, each of the methods were evaluated for their ability to detect enrichment of GO terms in the *Arabidopsis thaliana* and *Saccharomyces cerevisiae* gene expression datasets. These datasets were selected because of the well-developed GO term annotation of these two species genomes, and the consistent gene name annotation between datasets. The fewest terms enriched in the results of any of the methods were in the results of SOMs. Thus, to simplify visualisation, the GO term enrichment results from SOMs were excluded from the diagrams in Fig. 4 and can be found in Supplementary Tables S4 and S5.

**Figure 4.**
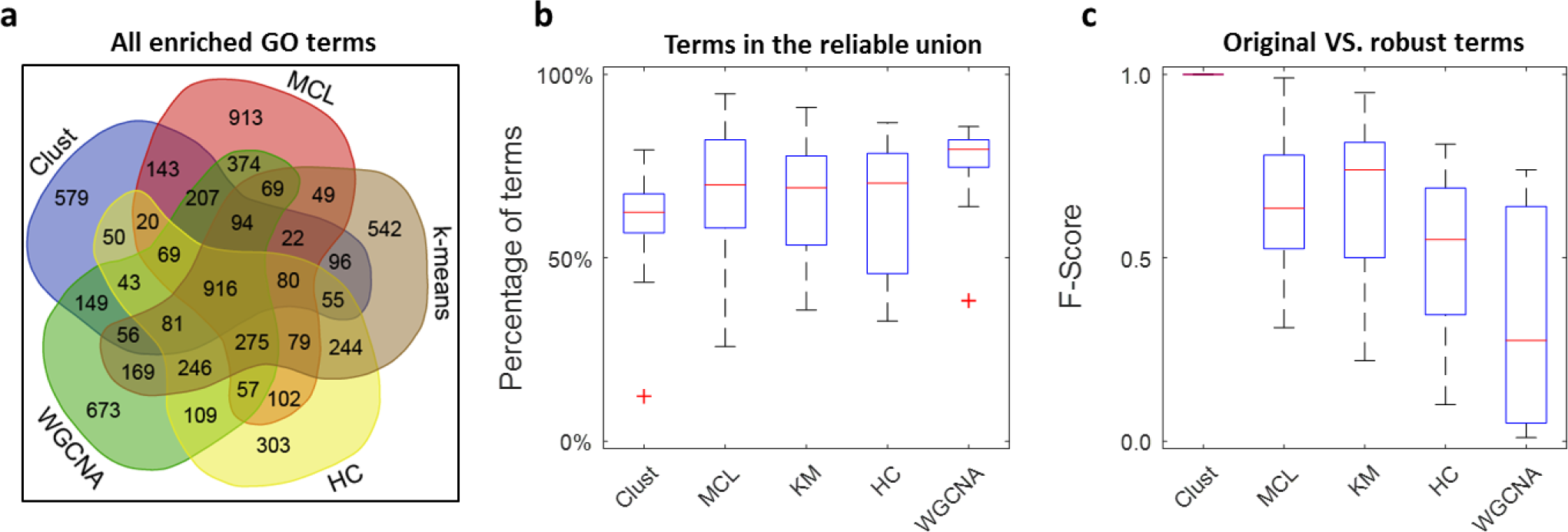
Evaluation of GO term enrichment in the results of the clustering methods. (**a**) Venn diagram demonstrating substantial differences in enriched GO terms between methods. The numbers on this diagram represent the number of GO terms detected as significantly enriched across the 20 selected datasets. The union of these sets includes 6864 terms, 3010 of which (44%) are exclusive to a single method. (**b**) The percentage of reliable GO terms (i.e. those detected by two or more methods) detected by each method. WGCNA is the only method which is significantly higher than clust (Supplementary Table S4). (**c**) F-scores quantifying the similarity between the set of GO terms detected as enriched in the original clusters and the set of GO terms detected as enriched in clusters after taking into account unreliably assigned genes (Supplementary Table S4).

Of the 6,864 significantly enriched GO terms detected by all methods (excluding SOMs), 3,854 (56%) were detected by two or more methods and 916 GO terms (13%) were detected by all methods (Fig. 4a). Given that the results were considerably different between methods, it is reasonable to assume that GO terms detected by two or more methods for a given input dataset are those that are more likely to be correct, however there was no significant difference between most of the methods in their ability to recover these GO terms (Fig. 4b and Supplementary Table S4 (A)). Thus, while *clust* only assigns 50% of genes to clusters, this reduction in gene content does not reduce the number of detected enriched functional terms compared to other methods.

To determine the effect of the inclusion or the exclusion of unreliably assigned genes on GO term enrichment, the clusters produced by the data partitioning methods we further analysed both with all of the unreliably assigned genes removed (the stringent set), and with all genes that fit within the boundaries of the cluster included (the expanded set) (Supplementary Figure S11). Those functional terms that remained significantly associated with a cluster irrespective of whether the stringent or expanded version of the cluster was analysed were deemed robust to unreliable gene content. On average for the data partitioning methods, between 10% and 80% of GO terms are not robust to this perturbation and are therefore unreliably assigned to clusters (Fig. 4c). Clusters produced by *clust* are discrete and therefore not affected by this unreliable assignment problem (Fig. 4c).

### Clust extracts clusters of co-expressed genes from multiple datasets simultaneously

The quantity of gene expression data that is deposited in public repositories is increasing rapidly. This is primarily due to a reduction in the costs of acquiring such datasets. These datasets come from a multitude of different species, have been generated using different technologies (microarrays and RNA-seq), and have different properties such as numbers of conditions, replicates, and missing values. *Clust* is designed to enable simultaneous cluster extraction from multiple such heterogeneous gene expression datasets (Supplementary Text S1). Irrespective of datatype or source species, *clust* extracts clusters of genes that are consistently co-expressed with each other in all of the given datasets.

To evaluate this feature of *clust*, ten combinations of *d* datasets (where *d* ∈ {2,3,4,5,6,7,8,9,10}) were selected at random from the ten yeast RNA-seq datasets (D91 to D100; Supplementary Table S1). The same experiment was performed over Arabidopsis datasets. To provide a comparison, the other methods were also applied to these combinations of datasets. However, as these methods are only applicable to a single dataset at a time, the only way to enable their simultaneous analysis was to concatenate them together prior to clustering (Supplementary Tables S6, S7, S8, and S9). As before, *clust* produces tighter clusters with lower within-cluster dispersion (lower MSE) (Fig. 5a & b) and guarantees no cluster profiles which overlap (JI = 0, Fig. 5c & d). Moreover, and as expected from a biological point of view, both the percentage of input genes that are included in the extracted clusters (PAG) and the number of generated clusters (*K*) decrease as more datasets are included as input to *clust* (Fig. 5e-h). This behaviour is expected because as the number of conditions increases, the less likely a group of genes are to be co-expressed under all conditions. For example, when all ten yeast RNA-seq datasets are provided as input to *clust*, only a single cluster of 56 genes is identified. Of these, 48 are components of the ribosome or participate in ribosome biogenesis (Supplementary Table S10). An analogous cluster of 50 genes (45 of which are ribosomal or involved in ribosome biogenesis) was obtained when all 10 Arabidopsis RNA-seq datasets were provided to *clust* (Supplementary Table S10). Of the other methods, MCL maintains relatively low MSE values over increasing numbers of datasets (d). In contrast, MSE values of the other methods increase when *d* increases (Fig. 5a-d). Moreover, MCL is the only method which shows a trend similar to *clust* in terms of decreasing values of PAG and *K* at higher *d* values (Fig. 5e-h). Nonetheless, the performance of clust is significantly better than MCL in terms of MSE and JI values at all *d* values (Fig. 5a-d).

**Figure 5.**
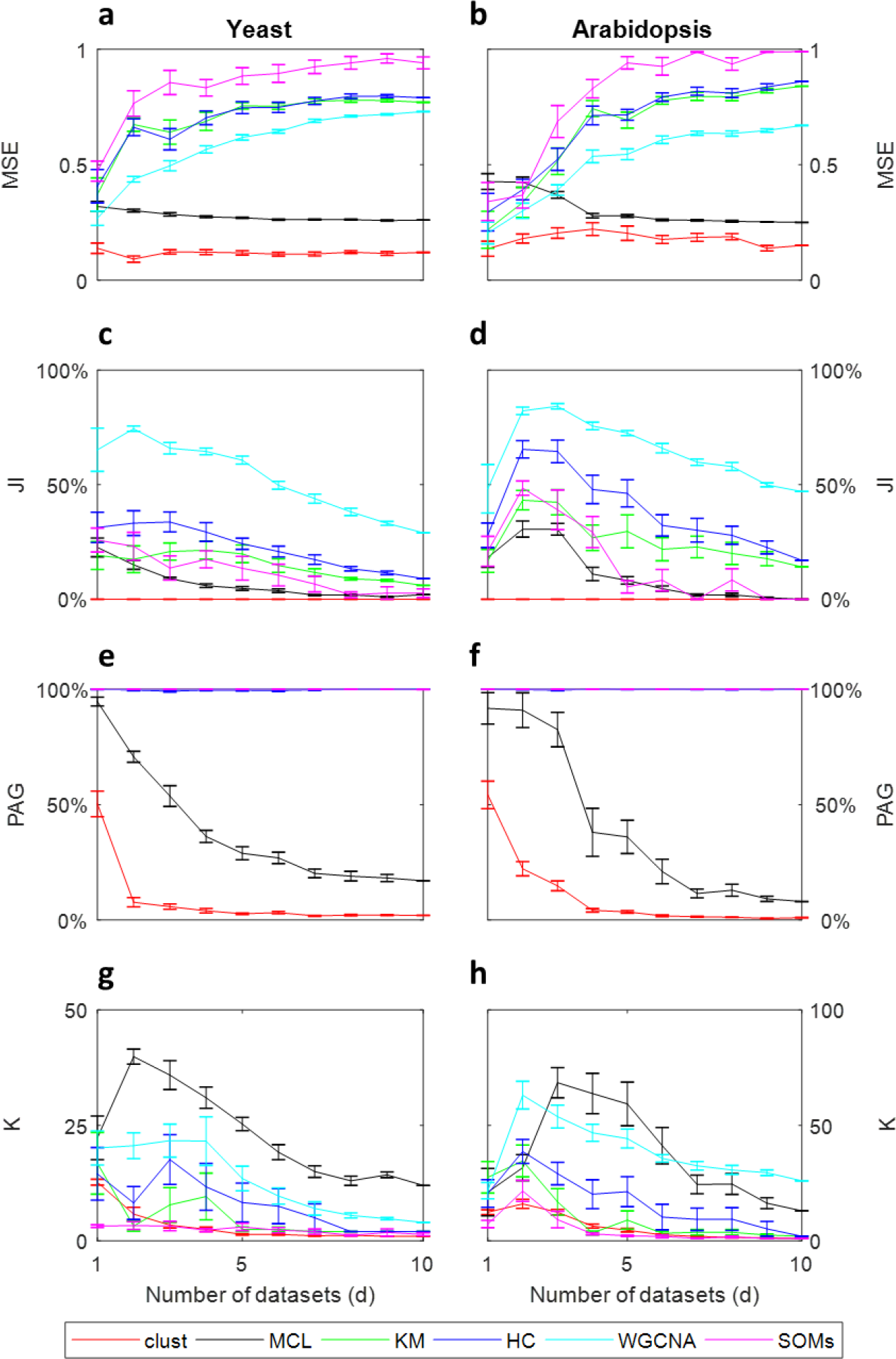
Evaluation of the performance of clustering methods when applied to multiple datasets. Each sub-plot shows the values of one performance metric measured for each method when applied on (d) different datasets simultaneously. The horizontal axes represent the numbers of datasets simultaneously clustered (d), the vertical axes represent the performance metrics’ values, and the error bars represent standard error values over 10 random repetitions. The datasets are from either yeast (**a**, **c**, **e**, and **g**) or arabidopsis (**b**, **d**, **f**, and **h**). The performance metrics are within-cluster dispersion measured by MSE (smaller values are better) (**a & b**), percentage of overlap amongst clusters measured by the JI index (smaller values are better) (**c & d**), percentage of genes assigned to clusters (PAG) (**e & f**), and number of clusters generated (K) (**g & h**).

## Discussion

Co-expression clustering is a routinely used step in data exploration for gene expression analysis. Here we show that the most commonly used methods for conducting co-expression analysis do not match the biological expectations of a co-expressed cluster of genes, producing clusters that are highly dispersed (high MSE values) and contain large proportions of genes that could be equally assigned to other clusters within the same clustering result (high JI values). Moreover, the methods behave inconsistently, with substantial differences in clustering performance attributable to differences in datatype or data quantity. We present *clust*, as a method designed to solve all of these problems. *Clust* was compared with five commonly used clustering methods (MCL, *k*-means, HC, WGCNA, and SOMs) by application to 100 different microarray and RNA-seq gene expression datasets from five model species. In contrast to the other tested methods, *clust* behaviour is consistent and is unaffected by species, datatype, number of genes, or number of conditions. Thus, *clust* performance is robust to increases in data quantity without sacrificing the quality of the results.

The most commonly conducted post-clustering analysis is to detect enrichment of functional terms within clustered sets of co-expressed genes. We show that conducting such analyses on clusters produced using the most commonly used methods for co-expressed gene clustering produces very different results (Fig. 4a) with between 10% and 80% of enriched functional terms being unreliably assigned to clusters (Fig. 4c). This observation has implications for the utility of downstream analysis conducted on these clusters. For example, putative regulatory relationships are often inferred by identifying regulatory genes that occur in clusters that are enriched for specific functional terms ^10,11,12^. Thus, unreliability of enriched functional term assignment likely contributes to the high false positive discovery rate in the discovery rate of regulatory interactions from co-expression data ^13^. As *clust* is designed to solve the problem of reliability of gene assignment to clusters, it does not suffer from such unreliability in enriched functional terms, and therefore represents a solution to this problem.

Finally, clust is designed to be able to extract clusters of co-expressed genes from multiple gene expression datasets, even if these datasets have different properties such as numbers of conditions or replicates. Such feature allows researchers who have multiple gene expression datasets that are all related to the biological problem in hand to analyse them simultaneously. That is, to extract the clusters of genes which are consistently co-expressed in each of these different datasets. Various consequences can be inferred from such analysis. For instance, it is more reliable to hypothesise that a group of genes are co-regulated by common regulator when they are consistently co-expressed over multiple datasets in contrast to being co-expressed in a single dataset only ^14,15,16,17,2^.

Taken together, this work reveals a mismatch between what researchers expect from gene expression clustering and the results that are produced by application of commonly used data partitioning methods to these data. The proposed *clust* method solves this problem, and the utility and performance characteristics of *clust* are demonstrated through comprehensive testing and comparison on real biological datasets from multiple different species. In addition to improved performance characteristics over competing methods, the ability of *clust* to handle multiple datasets simultaneously will enable individual gene expression datasets to be interpreted in the context of the large quantity of publicly available gene expression data. *Clust* is open source and freely available at https://github.com/BaselAbujamous/clust.

## Methods

### Overview of the clust cluster extraction method

*Clust* has a pipeline of steps that extract final optimised clusters of co-expressed genes from one or more gene expression datasets. A summary description of the algorithm is presented in this section and the full details are provided in Supplementary Text S1. A standalone Python implementation of *Clust* is available at https://github.com/BaselAbujamous/clust. In brief, *clust* begins by finding the general trends in the data and employs a number of base clustering methods (e.g. k-means clustering, hierarchical clustering, and self-organising maps) to produce initial sets of guide clusters. These initial clusters are provided as input to construct consensus “seed” clusters using Bi-CoPaM ^18^. All seed clusters are evaluated by the M-N distance metric ^19^ which considers both within-cluster dispersion and cluster size. Then, the set of non-overlapping clusters that minimise the M-N distance metric, that is, that minimise within-cluster dispersion while maximising cluster size are selected as elite seed clusters. The final step of the algorithm removes outliers from elite seed clusters using a Tukey filter, defines the cluster profile based on the range of expression values observed within the cluster, and then assigns all genes from the input dataset that fit within this cluster profile.

### Selection of 100 gene expression datasets

The 100 gene expression datasets were downloaded from the Gene Expression Omnibus (GEO) repository on 2^nd^ of July 2017 ^20^. For each one of the five model species, ten microarray datasets and ten RNA-seq datasets were downloaded. In all cases, the most recently published datasets for each of these species was selected, given that the dataset had at least 4 different conditions (time-points or treatments) and no more than 50 samples including replicates. RNA-seq datasets were chosen only if the resulting TPM, RPKM, FPKM, or CPM quantitation files were available from the GEO repository. Microarray datasets were a mix of both one-colour or two-colour microarrays. The complete list of the 100 datasets and their properties is available in Supplementary Table S1. The raw data files for the 100 datasets, the clustering from each method, and the analysis scripts are all publically available at the Zenodo repository with the doi 10.5281/zenodo.1169191.

### Implementation of clust and comparative methods

All methods, including *clust*, were run using their default parameters, which is the manner in which they are most commonly used. Running k-means, HC, and SOMs, requires pre-setting the number of clusters (*k*). Each of these methods was applied to the input data with k values ranging from 2 to 50 and the *k* value that minimised the Davies–Bouldin (DB) cluster validation index was chosen. The DB index is the most widely used and most frequently cited whole-partition cluster validation index ^9^.

k-means was run using the Python *sklearn.cluster* implementation. The Python *mcl* package was used to run MCL after generating networks of co-expressed genes using an intuitive Pearson’s correlation threshold of 0.8 ^21^. The Python *scipy.cluster.hierarchy* package was used to run HC clustering. The *blockwiseModules* module of the R *WGCNA* library was used to run WGCNA with the network type set to “signed”. The Python *sompy* package was used to run SOMs. The Python package *clust 1.2.0* was used to run *clust*. To demonstrate that the superior performance characteristics of *clust* were not due to use of the DB index, we also attempted to bias against our principle finding by choosing the cluster sets that minimised our evaluation criteria i.e that minimised MSE and the JI metrics (minimising 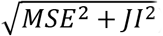). These additional results are analogous to those produced using DB index and are included in Supplementary Table S3 (B) and Supplementary Figure S2.

### Cluster dispersion metric (MSE)

The mean squared error (MSE) metric is used to measure within-cluster dispersion. If the cluster has *N* genes and the dataset has *D* dimensions, the MSE value for that cluster will be:

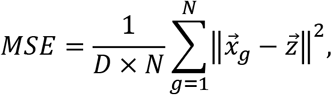

where 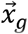 is a vector of the gene expression profile of the *g^th^* gene in this cluster, 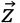 is a vector of the average expression profile of all genes in this cluster, and 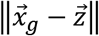 is the Euclidean distance between these two vectors. Note that the MSE value here is normalised by the number of genes in the cluster. When calculating the MSE value for a whole clustering result (a set of clusters), it is calculated as the weighted average of the MSE values of the each of the clusters, where the weight is the size (number of genes) in each of the clusters.

### Cluster similarity metric (JI)

The Jaccard Index (JI) metric is used to measure the similarity amongst the clusters in a clustering result ^22^. JI is calculated as the ratio between the number of “overlap genes” and the number of all genes in clusters. “Overlap genes” are those genes that are included in a cluster while their expression profiles also fit within the boundaries of at least one other cluster. The upper and the lower boundaries of a cluster at any given dimension (condition) are respectively calculated as the maximum and the minimum expression values of all genes in that cluster after trimming the most extreme 1% values at each point to reduce the effect of outliers.

### GO term enrichment analysis

The GO term annotations for *Arabidopsis thaliana* and *Saccharomyces cerevisiae* were downloaded from the Gene Ontology Consortium’s online repository at http://www.geneontology.org ^23,24^. Significantly enriched GO terms were taken as those that obtained an adjusted hypergeometric test p-value ≤ 0.001.

## Acknowledgements

BAJ and this work were supported by the Bill & Melinda Gates Foundation through award number OPP1129902. SK is a Royal Society University Research Fellow. Work in SKs lab is supported by the Royal Society, and the European Union’s Horizon 2020 research and innovation programme under grant agreement number 637765.

## Author contributions

BAJ and SK designed the study, analysed the data, and wrote the manuscript. BAJ developed the software and conducted the analysis.

